# Towards Principled Modeling Of Coronary Artery Calcium Scores With Zero-Inflated Regression

**DOI:** 10.1101/2024.08.06.606909

**Authors:** Avanti Shrikumar, Sina Fathieh, Gemma A. Figtree, Stuart M. Grieve

## Abstract

The Coronary Artery Calcium Score (CACS) is a widely-used measure of Coronary Artery Disease (CAD). The primary use of CACS is to identify subclinical CAD and estimate risk of future cardiovascular events such as acute myocardial infarction. The development of coronary atherosclerosis is well known to be accelerated in response to risk factors such as age, hypertension, hypercholeterolaemia and smoking. However, there is substantial variability in an individual’s susceptibility or resistance to CAD against these risk factors. Quantifiying the deviation from “expected” CACS provides an novel opportunity to inform multi-omic and similar unbiased discovery of new markers and mechanisms of CAD trained against CT imaging. Standard linear regression struggles to model CACS due to the high prevalence of zeros (which reflect the absence of measurable coronary plaque). Prior works have variously handled this by discarding measurements with CACS of zero or by binning the CACS into broad categorical groups. Such approaches discard meaningful data, motivating the need for a more principled approach to handling the data distribution. In this work, we explored zero-inflated regression as a possible approach to modeling CACS using a cohort of patients from the BioHEART-CT study, and devised metrics to validate performance. We identified zero-inflated negative binomial regression, zero-inflated gamma regression and zero-inflated lognormal regression as promising approaches for handling the distributional properties of CACS, where the best method to use can vary depending on the dataset considered. A key contribution of our work is to demonstrate how these models can also estimate the percentile of the observed CACS relative to the distribution that would be expected after controlling for the inputs, thereby avoiding the need for data-hungry binning-based approaches.

## 1 Introduction

Coronary Artery Disease (CAD) accounts for approximately half of all deaths due to cardiovascular disease and is a leading cause of death worldwide [1]. A significant fraction of patients who experience acute events or sudden death show no previous symptoms [2], motivating the use of the Coronary Artery Calcium Score (CACS) as a measure of cardiovascular risk. Coronary artery calcification results from the stabilization of atherosclerotic plaques, and can be detected using non-contrast Computed Tomography (CT). Conducted and quantified in a standardized manner, these images can generate a CACS that has been shown to be an effective predictor of the burden of coronary atherosclerosis and the subsequent risk of cardiovascular events [3].

There are many applications in which we may wish to model how an individual’s CACS relates to risk factors such as age, hypertension, hypercholeterolaemia and smoking. Beyond developing improved cardiovascular risk scores and discovering novel factors that may correlate with cardiovascular disease, a key application relevant to our study (BioHEART) was identifying groups of patients who do not follow a “typical” risk factor pattern (i.e. sub-groups of individuals whose CACS is unusually high or low relative to what would be predicted using known models of standard risk factors) in order to inform multi-omic and similar unbiased discovery of new markers and mechanisms of CAD. A challenge that arises when building such models is that the distributional properties of CACS are not well suited to naive least-squares regression, both due to the high prevalence of CAC scores of zero, as well as the fact that CAC scores can vary by orders of magnitude (illustrated in **Figure 1**). To date, there are no published guidelines advising researchers on how best to handle these distributional properties of CACS, which can lead researchers to avoid the modeling challenges by discarding meaningful information (such as by excluding patients with CACS of zero, or binning CACS into coarse-grained categorical groups).

**Figure 1:**
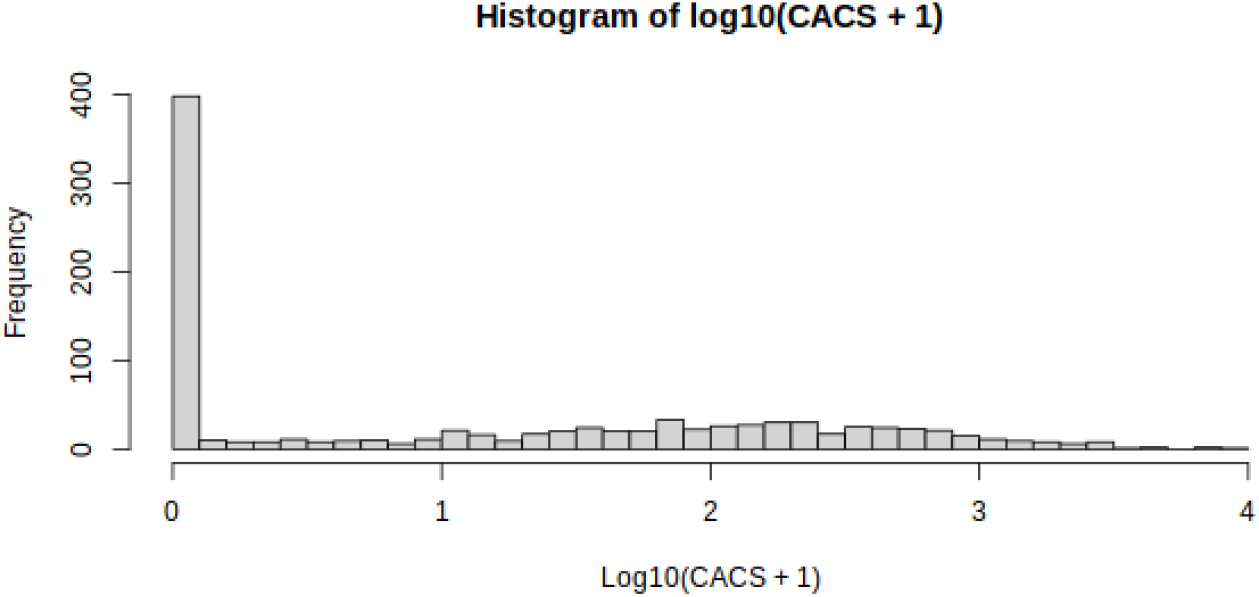
Histogram of CACS. Distribution generated based on 969 subjects from the BioHEART-CT study. 41% of CAC scores are zero.

This work is intended as a guide to more principled approaches to modeling CACS. We investigate various regression-based modeling approaches and identify zero-inflated negative binomial, zero-inflated gamma and zero-inflated lognormal regression as viable choices for modeling CACS, where the best method to use can vary depending on the dataset. We further show how, using such approaches, it is possible to derive principled estimates of how high or low a patient’s actual CACS is relative to the distribution that would be expected based on the patient’s inputs. Our recommendations could substantially improve the efficacy of studies involving CACS, as well as studies involving other measurements that might benefit from similar modeling approaches.

## 2 Background

### 2.1 What Do (Most) Regression Models Actually Predict?

While it is common to think of regression as predicting a single output value given provided input values, an often-overlooked feature of (most) regression models is that they predict the *distribution* that the output originates from after conditioning on the input values^1^. For example, in least-squares linear regression, the model can be viewed as implicitly predicting that the output originates from a Gaussian distribution whose mean is linearly dependent on the input values; minimizing the least-squares loss is equivalent to maximizing the likelihood of the data according to these Gaussian distributions.

Regression techniques other than least-squares linear regression make different assumptions about the output data distribution. For example, Poisson regression assumes that the output originates from a Poisson distribution whose mean depends on the input, and thus uses a different loss function than least-squares. **Figure 2** shows the predicted distributions from four families of regression models trained to predict the distribution of non-zero CAC scores at age=50 (see **Sec. 3.2** for more details).

**Figure 2:**
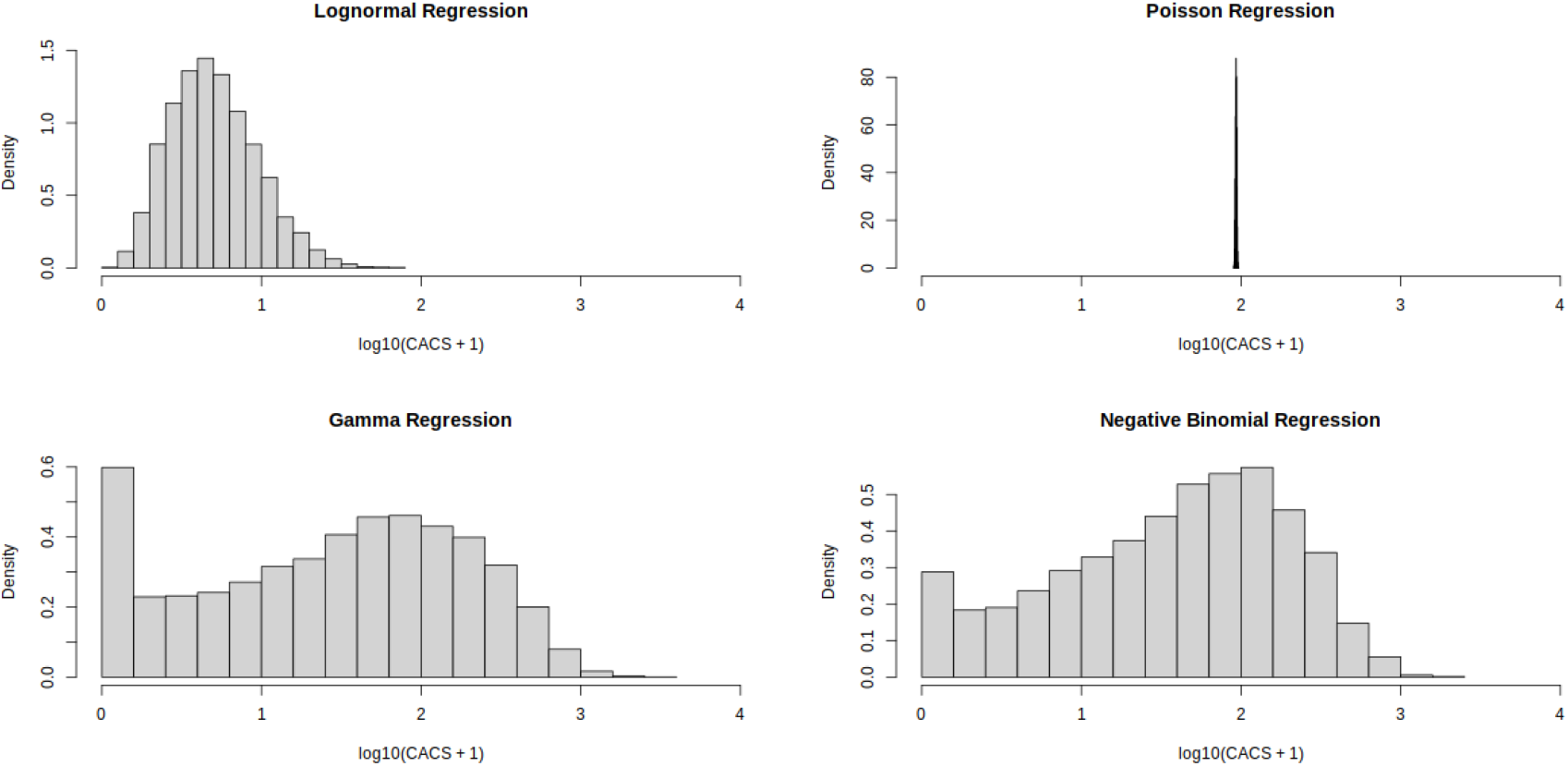
Distributions predicted for age=50 by different regression models. The models depicted here were trained only on non-zero CACS and are described more in Sec. 3.2. Poisson regression predicts a very narrow spread of CACS compared to lognormal regression, gamma regression, and negative binomial regression.

This feature of regression models allows us to compute the percentile of a subject’s observed CACS relative to the distribution of CACS that is predicted based on the subject’s input values. We refer to this type of input-dependent percentile as a “conditional” percentile. To our knowledge, we are the first to observe that CACS conditional percentiles can be computed directly from the predicted distributions of a regression model, without any need to group subjects into bins based on their input features (as was done in previous works [4]).

### 2.2 What is zero-inflated regression?

A zero-inflated distribution (such as zero-inflated Poisson or zero-inflated negative binomial) is one where a fraction of the values is predicted as zero, and the remaining fraction is drawn from another distribution. So, for example, a “zero-inflated negative binomial distribution” is one where a fraction of the values is drawn from the zero distribution and the remaining fraction is drawn from a negative binomial distribution (we write “drawn from the zero distribution” rather than just “zero” because a negative binomial distribution is also capable of producing zeros). In zero-inflated regression, the proportion of values predicted to originate from the zero distribution is allowed to vary depending on the inputs. To illustrate this, **Figure 3** shows the output distributions predicted at two different ages for a zero-inflated negative binomial regression model trained to predict CACS as a function of age.

**Figure 3:**
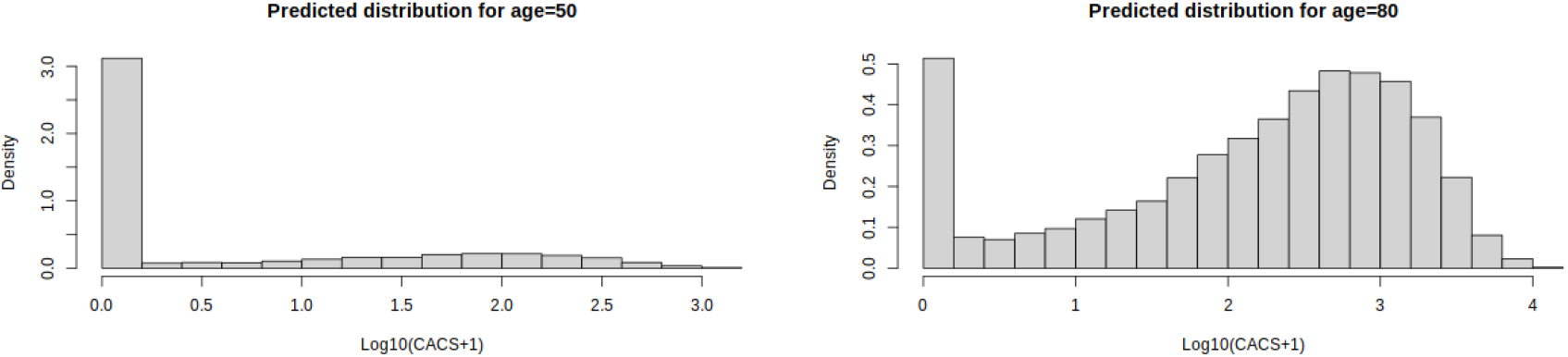
Distributions predicted for different ages by zero-inflated negative binomial regression. We see that the proportion of zeros predicted at age=50 is much higher than the proportion of zeros predicted at age=80.

Because the zero-inflation technique can work with a variety of distributions, we first focused on identifying those distributions that showed promise at modeling the non-zero CACS, and then combined them with zero-inflation to model the full CACS dataset.

## 3 Methods

### 3.1 Input datasets and evaluation tasks

Our full dataset consisted of a cohort of 969 subjects from the BioHEART-CT study, out of which 50% were assigned to a training set and 50% to a testing set. When evaluating which distributions were a good fit for modeling only the non-zero CACS, we limited our dataset to 571 subjects with nonzero CACS and again used a 50-50 split into training and testing sets.

Because our goal was to evaluate how well the distributional assumptions of different regression models fit with the CACS data, for most of the evaluations presented in this work we focused on the task of predicting the CACS as a function of age alone because it is difficult to visualize the relationship between the predicted percentiles and the input when the input is multidimensional. We also tested our methods on a task in which we predicted CACS as a function of 4 computed cardiovascular risk scores (FRS, ACCAHA, MESA and SCORE2) on a subset of 731 subjects for which the risk scores were available (the remaining subjects were either not in the age range for which the risk scores were designed or had missing data). For this secondary task, the computed risk scores were transformed using R’s `;orderNorm` function.

The distribution of CACS on the subset of 731 subjects with risk scores is notably different compared to the full set of 939 subjects: 45% of the 731 subjects have non-zero CACS, while among the remaining 238 for which risk scores were not available, only 27% have non-zero CACS. To investigate whether this distributional shift made a difference to which method performed the best, we also fit models to predict CACS based on age alone on this subset of 731 subjects.

### 3.2 Regression Models

In this section, we describe the regression models that were investigated in this work. We will use *o*_*i*_ denote the target output value for subject *i* (in this case, the subject’s CAC score), and 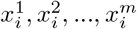 to denote the *m* input features for subject *i* (e.g. in the case of a model fit to the sole input of age, we have *m* = 1 and 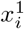 would be the age of subject *i*).

#### 3.2.1 Lognormal regression

A lognormal distribution is one whose log is normally distributed. Thus, lognormal regression can be performed on the non-zero CACS by running least-squares linear regression on log-transformed CACS, as the least-squares loss implicitly assumes that the outputs come from normal distributions. Specifically, in least squares linear regression, for each input feature *j* the model learns a parameter *β*_*j*_ such that observed output for subject *i* is assumed to originate from a Gaussian distribution with mean 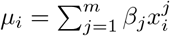. By identifying parameter values that minimize the least-squares loss, the model is maximizing the probability of the observed data under these Gaussian distributions. This is because the log probability that observation *o*_*i*_ originated from a Gaussian with mean *µ*_*i*_ is proportional to 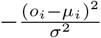, where *σ*^2^ is the variance of the Gaussian. If all observations are given equal weight, this is equivalent to assuming that all observations originate from Gaussian distributions with equal variance.

We performed lognormal regression by running R’s `;glm` function on the log transformed CACS (excluding CACS of zero). The resulting GLM stores the variance of the fit Gaussians as the “dispersion” parameter. To compute the percentile of a subject’s CACS relative to the distribution predicted by the fitted model, we used R’s `;pnorm` function with the quantile set to the subject’s log-transformed CACS, the mean set to the model’s predicted mean log-transformed CACS given the subject’s input features, and the standard deviation set to the square root of the dispersion parameter.

#### 3.2.2 Poisson regression

We fit a Poisson regression model to the dataset of non-zero CACS using R’s `;glm` function with `;family=“poisson”`. For each input feature *j*, the model learns a parameter *β*_*j*_ such that the observed output for subject *i* is assumed to originate from a Poisson distribution with mean 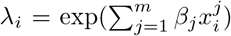. Because Poisson distributions are only intended for integer-valued data, for model fitting purposes we converted our CACS to integers by multiplying by 100 (the CACS were reported to 2 decimal places). To compute the percentile of a subject’s CACS relative to the distribution predicted by the Poisson regression model, we used R’s `;ppois` function with the quantile set to the subject’s CACS (multiplied by 100) and the lambda parameter set to the model’s predicted mean value for 100-times-CACS given the subject’s input features.

#### 3.2.3 Gamma regression

We fit a gamma regression model to the dataset of non-zero CACS using R’s `;glm` function with `;family=Gamma(link=“log”)`. For each input feature *j* the model learns a parameter *β*_*j*_ such that the observed output for subject *i* is assumed to originate from a gamma distribution with mean 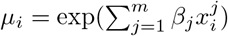.

Gamma distributions also include a “shape” parameter that impacts the variance of the predicted distributions. This shape parameter is assumed to be the same for all observations (analogous to how, in the case of least-squares linear regression, the variance of the predicted Gaussian distributions is assumed to be the same for all observations that have equal weight). As described in Ford (2022) [5], in the fitted glm this shape parameter is stored as the inverse of the “dispersion” parameter.

To compute the percentile of a subject’s CACS relative to the distribution predicted by the gamma regression model, we used R’s `;pgamma` function with the quantile set to the subject’s CACS, the shape parameter set to the inverse of the dispersion parameter, and the scale parameter set to the model’s predicted mean value for CACS divided by the shape (this relationship between the scale parameter, the shape parameter and the predicted mean is described in Ford (2022) [5]).

#### 3.2.4 Negative binomial regression

We fit a negative binomial regression model to the dataset of non-zero CACS using R’s `;glm.nb` function. For each input feature *j* the model learns a parameter *β*_*j*_ such that the observed output for subject *i* is assumed to originate from a negative binomial distribution with mean 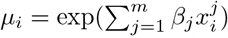. Because negative binomial distributions are only for integer valued data, for model fitting purposes we converted our CACS to integers by multiplying by 100.

The fitted negative binomial distributions also include a “size” parameter that impacts the variance of the predicted distributions. This parameter is assumed to be the same for all observations, and can be extracted from the fitted model (stored as the parameter “theta”). To compute the percentile of a subject’s CACS relative to the distribution predicted by the negative binomial regression model, we used R’s `;pnbinom` function with the quantile set to the subject’s CACS (multiplied by 100), the mean set to the model’s predicted mean value for 100-times-CACS, and the size parameter set to the theta value learned by the model.

#### 3.2.5 Zero-inflated lognormal regression

We considered zero-inflated lognormal regression in the context of modeling the full dataset of both zero and non-zero CACS. Because a lognormal distribution cannot have a value of zero, a “zero-inflated” lognormal model is equivalent to what is called a “hurdle” model in which two sub-models are fit completely separately and later combined. The two sub-models are a binary logistic regression trained to predict whether the CACS is nonzero, and lognormal regression trained solely on the non-zero CACS (using the approach described in **Sec. 3.2.1**).

Let 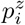 denote the probability predicted for subject *i* of drawing from the zero distribution, let *µ*_*i*_ denote the corresponding predicted mean (on a log scale) of the lognormal distribution, and let *σ* denote the standard deviation (on a log scale) obtained from the lognormal model as described in **Sec. 3.2.1**. The (maximum) percentile of a CACS of *o*_*i*_ is 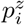 when *o*_*i*_ = 0, and 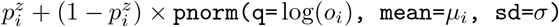 when *o*_*i*_ > 0.

#### 3.2.6 Zero-inflated gamma regression

We considered zero-inflated gamma regression in the context of modeling the full dataset of both zero and non-zero CACS. As with zero-inflated lognormal regression, because a gamma distribution cannot have a value of zero, we fit the “zero-inflated” gamma regression model using the hurdle model formulation in which a binary logistic regression model predicts whether the CACS is nonzero and standard gamma regression is performed solely on the non-zero CACS using the approach described in **Sec. 3.2.3**.

Let 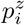 denote the probability predicted for subject *i* of drawing from the zero distribution, let *µ*_*i*_ denote the corresponding predicted mean (on a log scale) of the gamma distribution, and let *α* denote the model’s estimated shape parameter (computed as described in **Sec. 3.2.3**). Using the relationship that the scale *θ*_*i*_ of a gamma distribution is *θ*_*i*_ = *µ*_*i*_*/α*, the (maximum) percentile of a CACS of *o*_*i*_ is 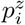 when 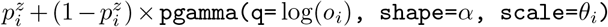 when *o*_*i*_ > 0.

#### 3.2.7 Zero-inflated negative binomial regression

We considered zero-inflated negative binomial regression in the context of modeling the full dataset of both zero and non-zero CACS. As before, to transform the CACS to integers for model fitting purposes, we multiplied the CACS by 100. We fit the model using the `;zeroinfl` function from the `;pscl` R package with `;dist=“negbin”` [6]. For each input feature *j* the model learns two parameters *β*_*j*_ and *γ*_*j*_ such that the observed output for subject *i* is assumed to come from a zero-inflated negative binomial distribution where the “zero” distribution is drawn from with probability 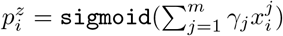, and the negative binomial part of the distribution has a mean of 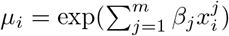 (as before, a “size” parameter is also learned for the negative binomial distribution and is stored as the constant “theta”).

Let 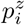 denote the probability predicted for subject *i* of drawing from the zero distribution, and let *µ*_*i*_ denote the predicted mean for the negative binomial part of the distribution. The (maximum) percentile of an observation *o*_*i*_ is then 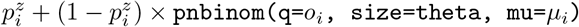.

### 3.3 A note on percentile ranges for zero-inflated models

For zero-inflated models it is possible to have a wide range in the percentiles at CACS=0, and thus care must be taken to consider both the maximum and minimum percentiles. The percentile formulas provided in the sections above are all for maximum percentiles. The minimum percentile of a CACS value of *o*_*i*_ is simply the maximum percentile of *o*_*i*_ − 0.01 (since the CACS is reported to two decimal places; if the CACS is multiplied by 100 as in the case of negative binomial regression, this would become *o*_*i*_ − 1). In other words, if we define our “maximum percentile” functions so as to return a percentile of zero for negative CACS, we can repurpose them to compute minimum percentiles by supplying a slightly smaller input. For the scatterplots involving zero-inflated models, the “percentile” of a CACS dataset refers to the median percentile (i.e. halfway between the minimum and maximum percentiles).

### 3.4 Scoring the probability of the observed data according to each model

One method of quantifying the fit of a model to a dataset is to compute the probability of observing the data according to the model. The probability of observing CACS within a range of a minimum of *a* to a maximum of *b* is simply the maximum conditional percentile of *b* minus the maximum conditional percentile of *a*. Because CACS is not precise to two decimal places, we compute the probability of observing a CACS within a range of +/-1 of the actual observed CACS. More precisely, if *g*(*o*) is a function that returns the maximum conditional percentile of observation *o*, we define the “probability” of observation *o* as *g*(*o* + 1) − *g*(`;max`(*o* − 1, 0.01)) if *o* > 0 and *g*(*o*) otherwise. The probabilities are converted to a log10 scale and averaged across subjects, with the results reported in corresponding figure titles in **Sec. 4**. Note that because probabilities are always less than 1, the log10 values are always negative, with more negative values indicating less probable observations. Additionally, because these scores quantify the probability of the specified range of CACS relative to the full range that could have been produced, the probabilities are relatively small. Finally, because the Poisson distribution is a poor fit to the distributional properties of CACS, due to machine precision limits the probabilities of some observations were computed as zero, resulting in log probabilities of negative infinity for Poisson regression.

### 3.5 Sanity checking the model calibration

Let us consider a model trained to predict the distribution of CACS for a given age. If the percentiles we assign using the model’s predicted output distribution are accurate, it means that if we were to hypothetically observe enough subjects with a given age, we would find (by definition of a percentile) that exactly *p*% of subjects with that age were assigned percentile values that were *<*= *p*%. In practice, the number of subjects in our dataset that have the same age may be limited, so we cannot reliably perform this check for individual ages. However, if this property that *p*% of subjects receive a percentile value *<*= *p*% is true in expectation for individual ages, then it holds true in expectation if we were to aggregate across all ages. In other words, if we were to use a model to predict conditional percentiles for all subjects in a dataset, and we were to plot the fraction *f* (*p*) of all subjects that received a conditional percentile *<*= *p*%, then we should expect that (for a model that predicts output distributions accurately) the plot of *f* (*p*) versus *p* would look like the *x* = *y* line.

We refer to these plots as calibration plots. Mathematically, they are equivalent to the Cumulative Distribution Function (CDF) of the conditional percentiles predicted by a model. Although these plots cannot tell us how accurate the predicted percentiles are for individual regions of the input space, they can tell us whether the proportions of predicted percentiles resemble the ideal proportions when looking across the full space of inputs.

In the case of the zero-inflated models that provide a range of percentile values, to generate these calibration plots we randomly sample a percentile value uniformly within the predicted percentile range for each observation, and repeat this 100 times.

## 4 Results

### 4.1 Evaluation on non-zero CACS

Our first goal was to identify methods that showed promise at modeling the distributional properties of the non-zero CACS. To this effect, we explored lognormal regression, Poisson regression, gamma regression and negative binomial regression. We chose the simple task of predicting CACS as a function of age, as this allowed us to visualize how the quantiles of the predicted CACS distributions changed with age. The results for the full set of 969 subjects (out of which 571 had nonzero CACS) are shown in **Figure 4**. We find that Poisson regression predicts distributions with very low variance, demonstrating that it is a poor fit for the distributional properties of CACS, and thus we excluded it from subsequent evaluations.

**Figure 4:**
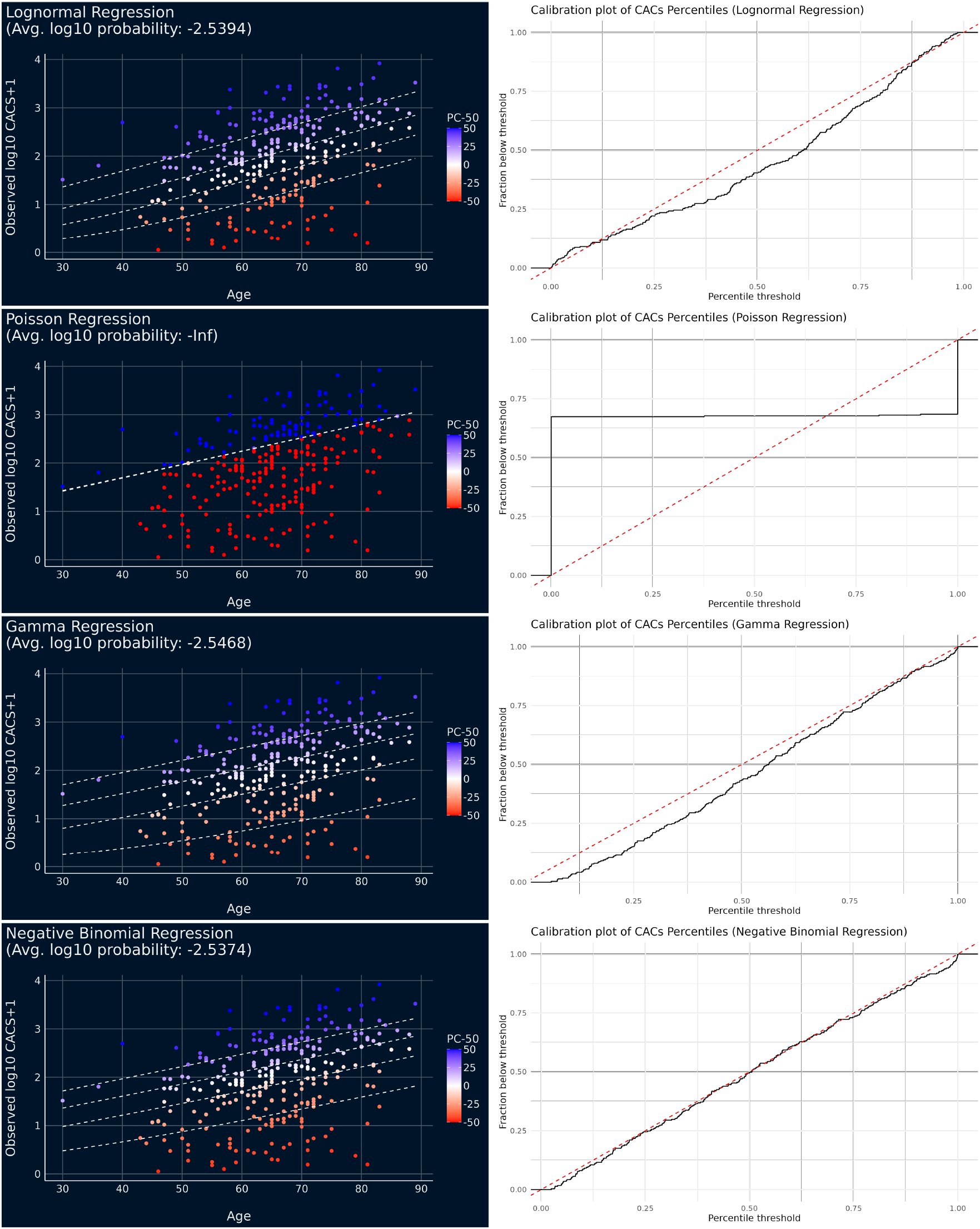
Percentiles computed on the testing set of the full 969-subject dataset by regression models trained to predict non-zero CACS as a function of age. The models depicted here are described in Sec. 3.2. Left panels: points are colored by the percentiles predicted by the model; the label “PC-50” on the scale refers to the percentile assigned by the model’s predicted distribution minus 50. From bottom to top, the dashed lines show the quantiles corresponding to percentiles of 20%, 40%, 60% and 80%. Right panels: calibration plot showing the fraction of observations that received a conditional percentile below a given threshold. Poisson regression predicts a very narrow spread of CAC scores compared to the other methods. The calibration plot of negative binomial regression is very close to the x=y line on this dataset.

We also find that, on the dataset that draws from the full set of 969 subjects, the calibration plot for negative binomial regression is almost perfectly in line with the x=y line. However, when we perform the identical evaluation using the subset of 731 subjects (**Figure 5**) for which the 4 cardiovascular risk scores could be computed (399 had nonzero CACS), we find that the calibration plot for negative binomial regression tends to lie slightly above the x=y line, suggesting a systematic underestimation of percentiles (as the actual fraction of subjects below a given percentile threshold is larger than the threshold). This underscores how the best modeling approach is sensitive to the CACS dataset under consideration.

**Figure 5:**
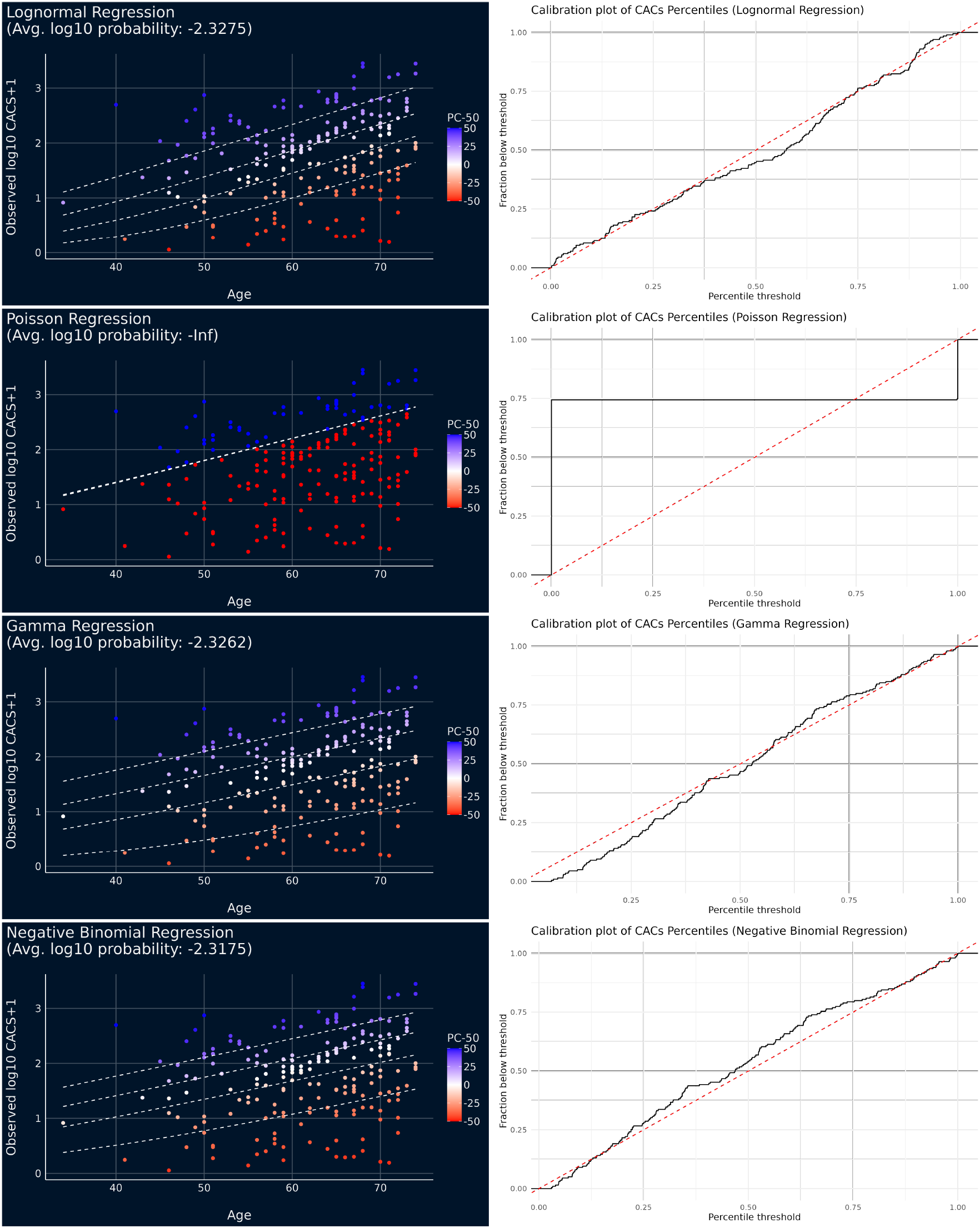
Percentiles computed on the testing set of the 731-subject dataset (containing only those subjects with 4 computed cardiovascular risk scores) by regression models trained to predict non-zero CACS as a function of age. Left panels: points are colored by the percentiles predicted by the model; the label “PC-50” on the scale refers to the percentile assigned by the model’s predicted distribution minus 50. From bottom to top, the dashed lines show the quantiles corresponding to percentiles of 20%, 40%, 60% and 80%. Right panels: calibration plot showing the fraction of observations that received a conditional percentile below a given threshold. Here, negative binomial regression does not appear to lie along the x=y line as neatly as in **Figure 4**

### 4.2 Evaluation on full CACS dataset

For evaluating the model performance on the dataset of both zero and non-zero CACS, we began with the simple task of predicting CACS as a function of age. The results, shown in **Figure 6**, illustrate how zero-inflation allows the predicted distributions to include large fractions of zeros, particularly at lower ages. The corresponding calibration plots (calculated as described in **Sec. 3.5**) show that the overall proportions of the predicted conditional percentiles have a slight bias towards underestimated percentiles in all three regression methods (as shown by the plots tending to lie above the x=y line). However, as demonstrated in **Figure 7**, this trend of overestimated percentiles was not present among the subset of 731 subjects for which risk scores could be computed, indicating that distributional shifts in the input dataset can impact how well zero-inflation is able to model the dataset.

**Figure 6:**
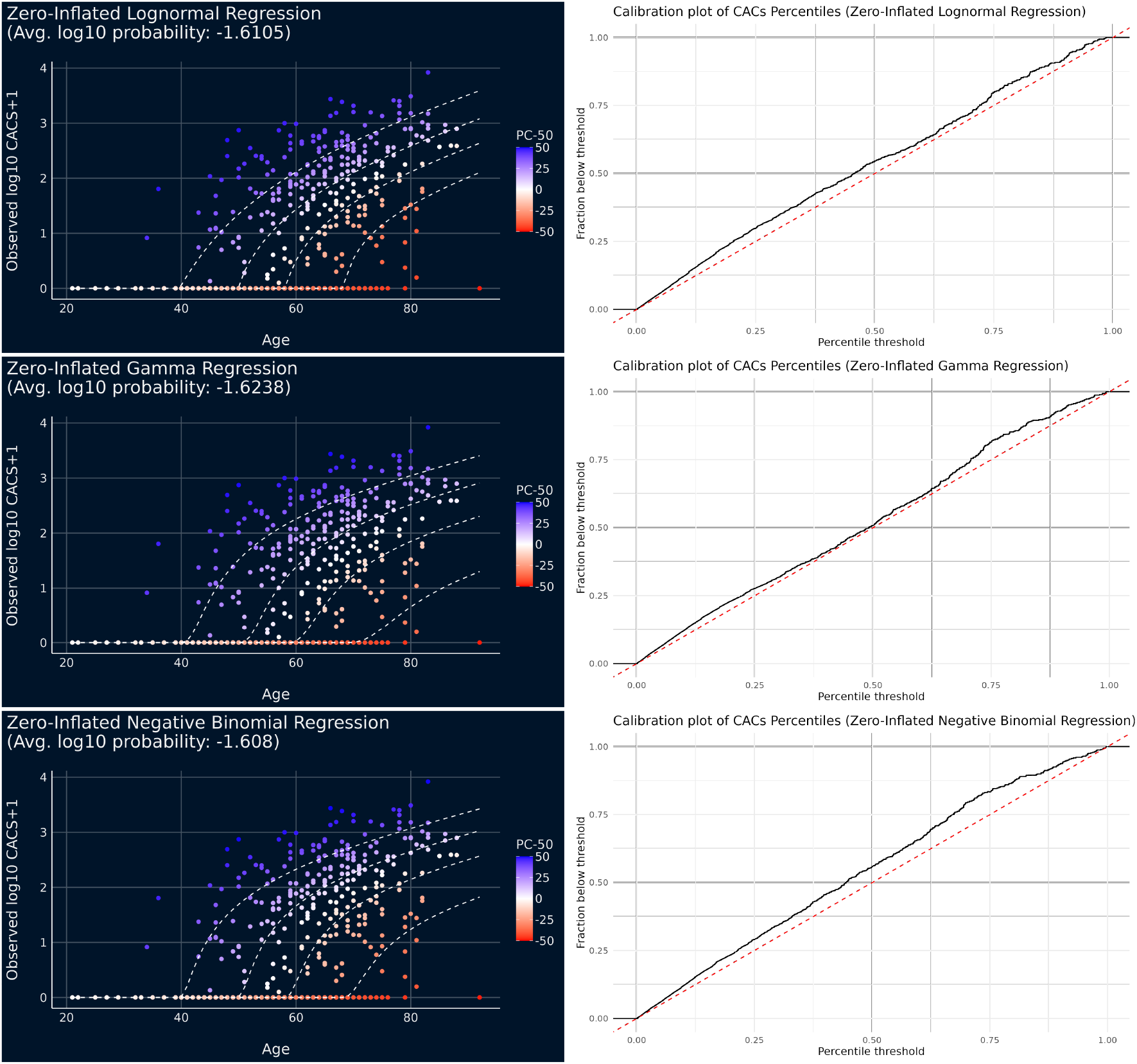
Percentiles computed on the testing set of the full 969-subject dataset by regression models trained to predict both zero and non-zero CACS as a function of age. Left panels: points are colored by the percentiles predicted by the model; the label “PC-50” on the scale refers to the percentile assigned by the model’s predicted distribution minus 50. From bottom to top, the dashed lines show the quantiles corresponding to percentiles of 20%, 40%, 60% and 80%. Right panels: calibration plot showing the fraction of observations that received a conditional percentile below a given threshold.

**Figure 7:**
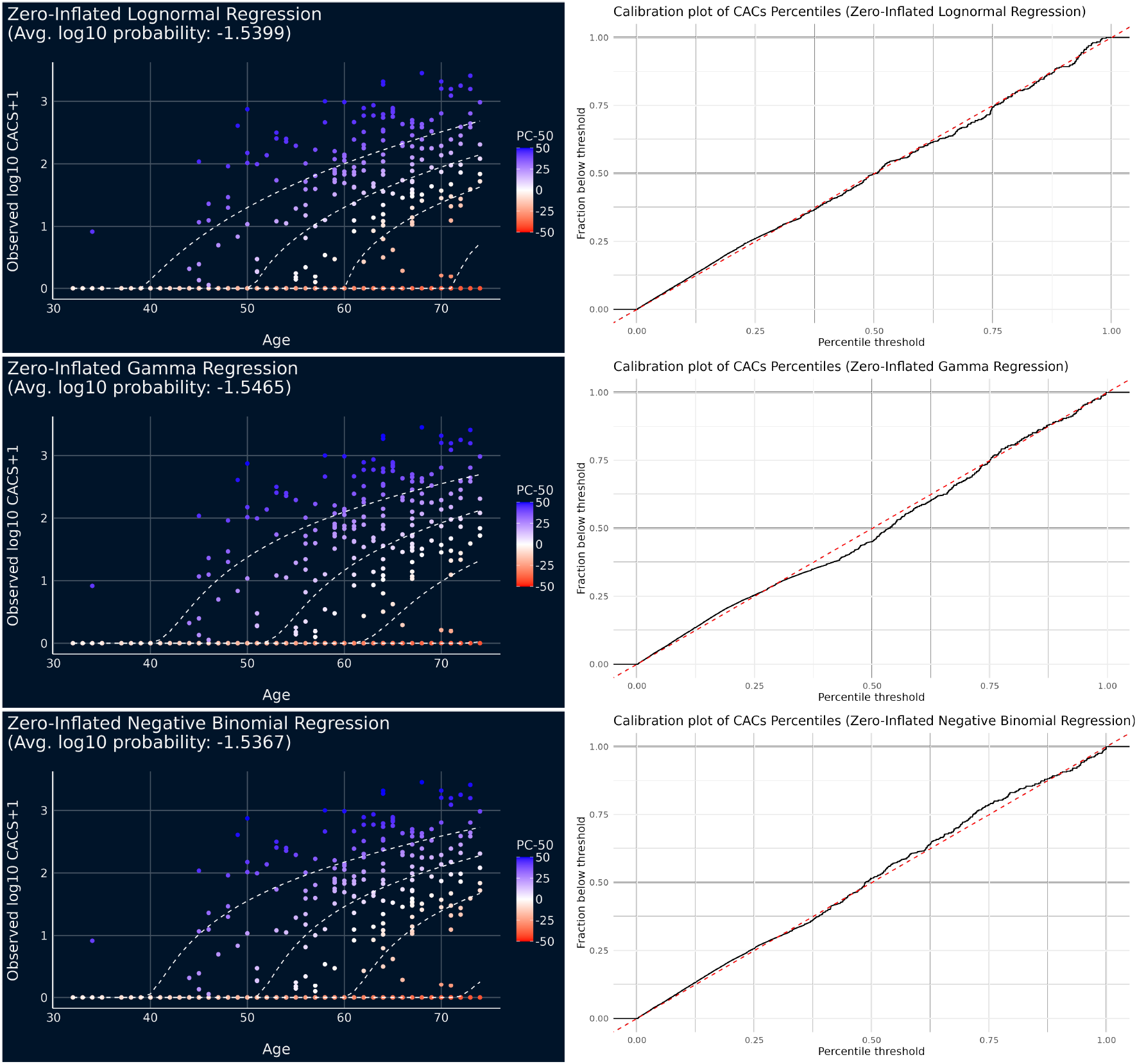
Percentiles computed on the testing set of the 731-subject dataset (containing only those subjects with 4 computed cardiovascular risk scores) by regression models trained to predict both zero and non-zero CACS as a function of age. Left panels: points are colored by the percentiles predicted by the model; the label “PC-50” on the scale refers to the percentile assigned by the model’s predicted distribution minus 50. From bottom to top, the dashed lines show the quantiles corresponding to percentiles of 20%, 40%, 60% and 80%. Right panels: calibration plot showing the fraction of observations that received a conditional percentile below a given threshold.

Finally, we evaluated our models on the more complex task in which the inputs were 4 computed cardiovascular risk scores (FRS, ACCAHA, MESA and SCORE2). The results are shown in **Figure. 8**. Due to the multidimensional nature of the input space, on the x-axis we show the mean predicted CACS, and we omit the dashed quantile lines as the quantiles are not a monotonic function of the mean predicted CACS (this is because zero-inflated regression involves two predicted parameters: the mean of the nonzero distribution as well as the proportion of observations drawn from the zero distribution). On this task, the calibration plots show that the overall proportions of the predicted conditional percentiles are in line with the ideal indicated by the x=y line.

**Figure 8:**
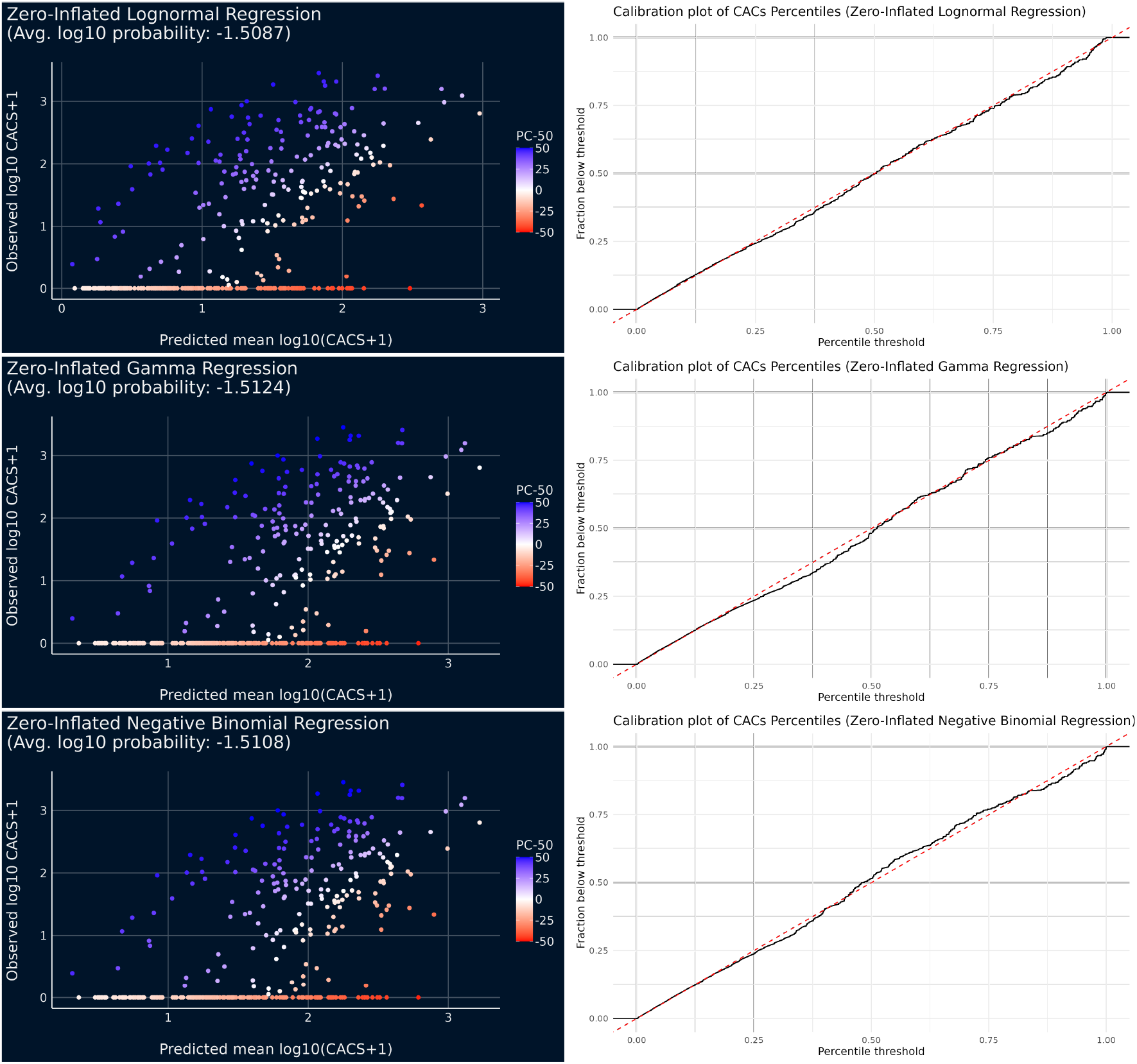
Percentiles computed on the testing set by models trained to predict CACS as a function of 4 computed cardiovascular risk scores. Left panels: points are colored by the percentile predicted by the model; the label “PC-50” on the scale refers to the percentile assigned by the model’s predicted distribution minus 50. Due to the complexity of visualizing a multidimensional input space, the x-axis is the model’s predicted mean CACS for each observation. Because zero-inflated models have parameters for both predicting the probability of zero and for predicting the mean value if non-zero, the percentiles do not monotonically increase with the model’s predicted mean. Right panels: calibration plot showing the fraction of observations that received a conditional percentile below a given threshold.

## 5 Discussion

In this work, we identified zero-inflated negative binomial regression, zero-inflated gamma regression and zero-inflated lognormal regression as promising approaches to modeling the distributional properties of CACS, where the best method to use can vary depending on the dataset. We found that Poisson regression is a poor fit for modeling CACS because it greatly underestimates the variance of output distributions. This is consistent with the fact lognormal, gamma and negative binomial distributions all contain a parameter that can scale the variance of the predicted distributions, while in Poisson distributions the variance is forced to be equal to the mean. While this feature of a Poisson distribution might have gone unnoticed in the simpler context of building a model to predict cardiovascular risk, it made a crucial difference in our more nuanced application of estimating deviation from median predicted risk via a conditional percentile.

While our findings focused on CACS, the broader issue of how to model distributions with a large fraction of zeros appears in other medical imaging contexts. For example, late gadolinium enhancement in the heart produces a distribution with a large fraction of zeros [7] that may be modeled well using zero-inflated regression.

To our knowledge, this work is the first to highlight that input-conditional percentiles for CACS can be derived directly from the distributions that are implicitly learned by regression models trained on CACS. This proposed approach circumvents the need for deriving such percentiles from data-hungry binning-based approaches and has the potential to boost the power of studies that rely on such percentiles for downstream analysis.

The calibration test proposed in this work can be applied to evaluate the predicted distributions of any regression model. However, it should be noted that because these calibration plots are computed across the full space of inputs, they can conceal issues wherein a model systematically overestimates percentiles for one region of the input space and underestimates percentiles in another region of the input space. Future work could explore additional metrics to evaluate the quality of a model’s predicted distributions.

Although this work focused on models in which the parameters of the predicted distributions were linearly dependent on the inputs, the overall finding generalizes to more complex models. For example, if a researcher wished to explore using an artificial neural network to predict the CAC scores, they would still be advised to consider designing their network predict one of the zero-inflated distributions suggested in this work as the output distribution (as opposed to defaulting to “least squares” optimization, which would implicitly predicting Gaussian distributions). We hope this work will inspire researchers to build on our findings to devise even more appropriate distributions and evaluation metrics to handle the properties of CACS and other scores.

## Acknowledgments

We thank Matthew Shu for preparing the input dataset of cardiovascular risk scores used to create **Figure 8**. We thank Arnaud Mazier for feedback on the presented work.

## Footnotes

1 We write “most” because this is only true of regression models that fall into the “maximum likelihood” framework, which covers most commonly-used models; there are some exceptions, such as quasi-Poisson regression

## Notes

### Competing Interest Statement

The authors have declared no competing interest.

